# Novel keto-alkyl-pyridinium antifungal molecules active in models of in vivo *Candida albicans* vascular catheter infection and ex vivo *Candida auris* skin colonization

**DOI:** 10.1101/2023.01.19.524835

**Authors:** Sarah R. Beattie, Taiwo Esan, Robert Zarnowski, Emily Eix, Jeniel E. Nett, David R. Andes, Timothy Hagen, Damian J. Krysan

## Abstract

New antifungal therapies are needed for both systemic, invasive infections as well as superficial infections of mucosal and skin surfaces as well as biofilms associated with medical devices. The resistance of biofilm and biofilm-like growth phases of fungi contributes to the poor efficacy of systemic therapies to non-systemic infections. Here, we describe the identification and characterization of a novel keto-alkyl-pyridinium scaffold with broad spectrum activity (2-16 µg/mL) against medically important yeasts and moulds, including clinical isolates resistant to azoles and/or echinocandins. Furthermore, these keto-alkyl-pyridinium agents retain substantial activity against biofilm phase yeast and have direct activity against hyphal *A. fumigatus*. Although their toxicity precludes use in systemic infections, we found that the keto-alkyl-pyridinium molecules reduce *C. albicans* fungal burden in a rat model of vascular catheter infection and reduce *Candida auris* colonization in a porcine ex vivo model. These initial pre-clinical data suggest that molecules of this class may warrant further study and development.

## Introduction

The effective treatment of human fungal infections is reliant on a relatively small set of structurally and mechanistically distinct drugs: azoles, polyenes, and echinocandin/triterpenoid glucan synthase inhibitors (1). As has been extensively discussed in the literature, this limited antifungal pharmacopeia must be expanded to meet the challenging clinical needs of patients in the 21^st^ century (2). Two specific examples where current antifungal therapies are relatively ineffective are: 1) infections involving biofilm on medical devices and mucosal surfaces (3) and 2) decolonization of *Candida auris* from skin (4). As with antibacterial agents, fungal biofilms are highly resistant to the currently used antifungal drugs (3). Consequently, removal of *Candida* infected medical devices such as vascular catheters and prosthetic joints is the standard of care (5), whereas devices infected with some species of bacteria are frequently salvaged with antibiotic therapy alone (6).

*C. auris* is an emerging fungal pathogen with many unusual characteristics compared to other *Candida* species (3). For example, it is readily transmitted from one patient to another in the hospital setting (7). This is very uncommon for other fungal pathogens and is likely due to its ability to persistently colonize patient’s skin as well as the inanimate surfaces in hospitals and long-term care facilities. Standard biocides used to decolonize microbes (e.g., methicillin-resistant *Staphylococcus aureus*) from patient’s skin such as chlorhexidine have relatively poor activity against *C. auris* (8). Furthermore, *C. auris* is frequently resistant to at least one of the three major classes of antifungal drugs (9) and pan-resistant isolates have been reported with increasing frequency (10). As such, *C. auris* is a pernicious problem that can result in therapeutically intractable infections.

One approach to treating surface-associated fungal infections such as biofilms and skin colonization is to identify molecules that are mechanistically distinct from those used to treat systemic infections and are specifically developed for that purpose. Such molecules could be imagined to display the general properties of traditional antiseptics but with activity optimized for specific organisms or improved safety profile. Here, we describe a set of novel N-keto alkyl pyridinium ions with broad spectrum antifungal activity against both planktonic and biofilm phase fungi.

Pyridinium-based anti-septic/anti-infective molecules have been studied and developed for topical and, in some instances, possible systemic use. Molecules in this class have been explored for antifungal applications (11, 12). For example, the long chain N-alkylated pyridinium cetylpyridinium chloride, a component of commercial mouthwashes, has in vitro activity against *Candida* spp with minimum inhibitory concentrations (MIC) of 2-6 µg/mL in modified CLSI conditions (13). A recent repurposing screen found that the anti-parasitic pyridinium-class drug pyrvinium pamoate is active against *C. auris* (MIC 2 µg/mL, ref. 14). Bis(pyridinium) molecules with variously sized alkyl or aryl linkers have also been shown to have antifungal activity (11, 15). One study showed that the bis(pyridinium) compounds induced less red blood cell lysis than similar bis(quaternary-ammonium) compounds (15).

The series characterized in this report are mono-pyridinium ions with aryl-alkyl ketone substituents at the N atom of the pyridinium ring (Fig. 1A). The molecules are comparable in activity to the commercially used, cetylpyridinium chloride and pyrvinium pamoate and have activity in an in vivo rat model of *C. albicans* vascular catheter infection as well as in an ex vivo model of *C. auris* skin colonization.

**Figure 1.**
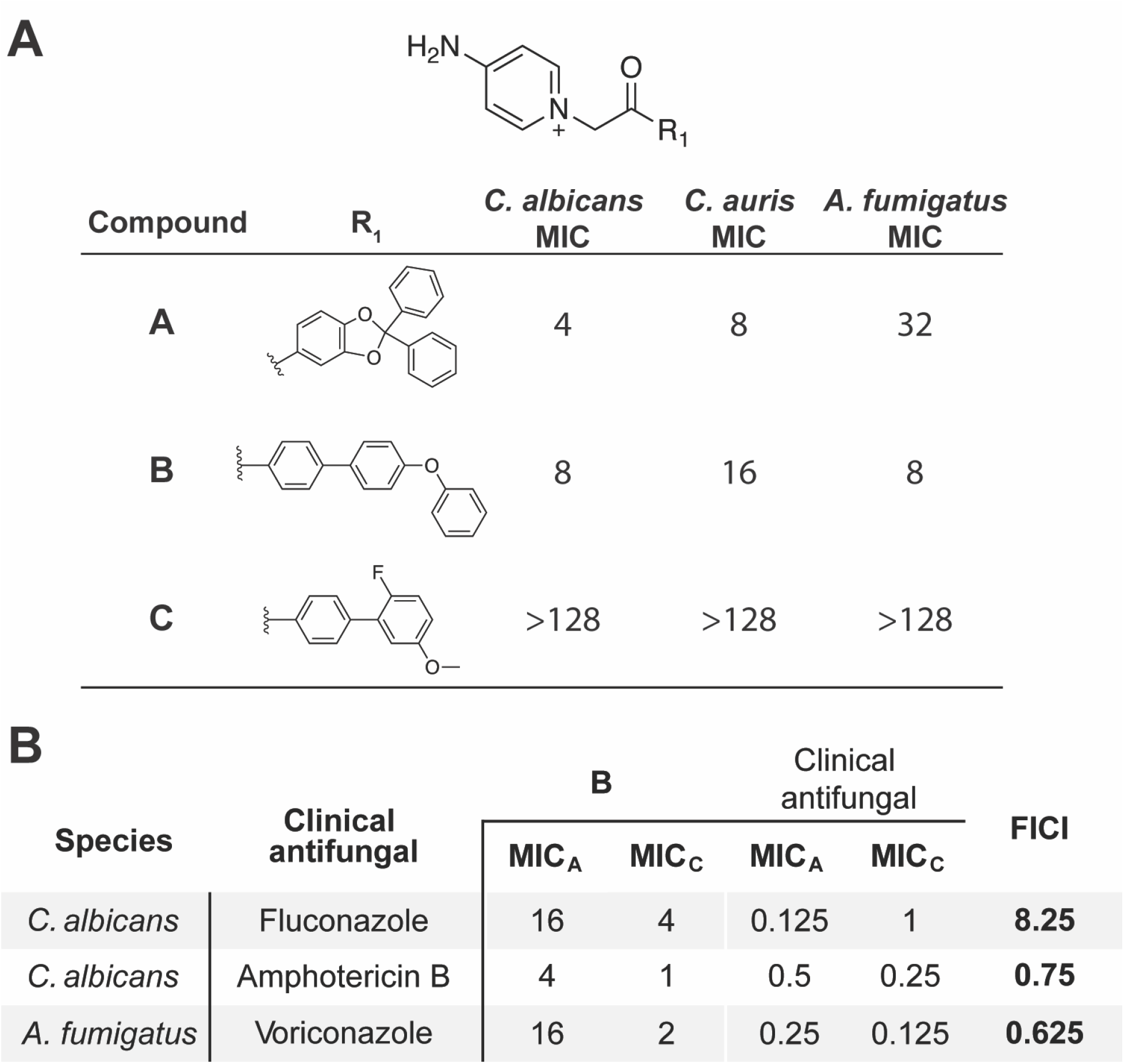
Ketoalkylpyridinium compounds with aryl substitutions have tunable antifungal activity. **A**. The pharmacore and R_1_ substitutions for compounds A, B and C with minimum inhibitory concentrations (MIC; µg/mL) for *C. albicans* SC5314, *C. auris* 0381, and *A. fumigatus* CEA10. **B**. Fractional inhibitory concentration indices (FICI) of B with clinical antifungals. FICIs were calculated using the MIC (μg/mL) of each drug alone (MIC_A_) and in combination (MIC_C_) based on at least two independent checkboard assays performed on separate days. Interactions are defined as: 0.5 ≥ FICI = synergy; 0.5 < FICI > 2 = additive; 2 < FICI > 4 = indifferent; FICI ≥ 4 = antagonism.

## Results

### Aryl methyl keto-*N*-alkylated pyridinium molecules have broad spectrum antifungal activity

A set of small molecule libraries were screened for compounds against *Aspergillus fumigatus* using recently developed high throughput screening assay applicable for this filamentous mould (16). A molecule with the aryl methyl keto-N-alkylated pyridinium (**k**eto-**a**lkyl **p**yridinium, KAP) scaffold (Fig. 1A) was identified a hit in the screen. A set of analogs of this initial hit were prepared and tested, leading to two derivatives with similar activity profiles (compounds **A** and **B**, Fig. 1A); **C** was used a control as it is inactive but has a very similar chemical structure compared to the active derivatives. Although >30 KAP derivatives were prepared, no clear structure-activity relationships (SAR) could be identified to explain the variation in antifungal properties of this series (see Fig. S1 for general synthesis approach). Pyridinium cations are widely believed to target membranes as part of their mechanism of action and a non-protein target could explain this difficult-to-interpret SAR (13, 17). Although other potential targets have also been proposed, a biochemically or genetically confirmed alternative target to the membrane has not been reported (12, 14). It is, however, interesting that small changes in substitution patterns in comparing A/B to C (Fig. 1A) lead to dramatic changes in antifungal activity without clearly identifiable changes in physicochemical properties. These observations suggest that the molecules have structurally specific interactions with a target and imply that interaction with membranes is not solely due to their amphipathic properties.

The scope of the antifungal activity for compounds A and B was explored by CLSI methods for a range of fungal pathogens including yeasts and moulds (Table 1). In general, A/B are more consistently active against yeasts compared to moulds (*A. fumigatus, A. terreus*, and *F. oxysporum*). Although the MIC for some isolates of *A. fumigatus* was similar to those for yeast, significant strain-to-strain variability in activity toward *Aspergillus* was observed. We also tested A/B against two fluconazole-resistant *C. albicans* strains (TWO15/17, ref. 17) that have increased expression of efflux pumps. We no difference in susceptibility compared to the reference strain SC5314 (Table 1). Furthermore, compounds A and B were both active against drug susceptible (0381) and multi-drug resistant (0390) *C. auris* isolates (Table 1).

**Table 1.** Minimum inhibitory concentration of KAP compounds against medically important fungi and bacteria. Minimum inhibitory concentrations (MIC; µg/mL) were measured using standard CLSI broth microdilution assays. Reported MICs are representative of at least two independent assays performed on different days. NT = not tested.

| Species | Strain | A MIC | B MIC |
| --- | --- | --- | --- |
| *A. fumigatus* | CEA10 | 32 | 8 |
|  | AF293 | 2 | 1 |
|  | SPF98 | 64 | 16 |
|  | Cas<sup>R</sup> | 32 | 16 |
|  | CFF S014 | 8 | NT |
|  | CFF S008 | 64 | NT |
|  | CFF S007 | 64 | NT |
|  | CFF S001 | 64 | NT |
|  | CFF S029 | 4 | NT |
|  | CFF S003 | 4 | NT |
| *A. terreus* | CFF S010 | 64 | NT |
|  | CFF S023 | 2-4 | NT |
| *C. albicans* | SC5314 | 4 | 8 |
|  | TWO #15 | 4 | 4 |
|  | TWO #17 | 4 | 4 |
| *C. glabrata* | KK2001 | 4 | 1 |
| *C. auris* | 0381 | 8 | 16 |
|  | 0390 | 2 | 4 |
| *C. neoformans* | H99 | 4 | 2 |
|  | DUMC 118.00 | 4 | 2 |
|  | DUMC 122.01 | 4 | 4 |
| *F. oxysporum* | DUMC Fo | 16 | 16 |
| *P. aeruginosa* | PAO1 | 64 | 32 |
| *S. aureus* | Newman | 32 | 16 |

Since pyridinium cations such as cetylpyridinium chloride also have activity against bacteria, we tested compounds A and B against the Gram-positive bacteria, *Staphylococcus aureus*, and the Gram-negative bacteria, *Pseudomonas aeruginosa*. Compared to cetylpyridinium choloride, compounds A and B were ∼32-fold less active against *S aureus* but were more active against *P. aeruginosa* (32 µg/mL vs 250 µg/mL reported in ref 17). In contrast, the activity of A/B were within 2-fold of the MICs reported for cetylpyridinium chloride against *C. albicans* and pyrvinium pamoate against *C. auris*.

To determine if compound interacted with clinically used antifungal drugs, we performed checkerboard assays and determined fractional inhibitory concentration indices. Compound B showed additive interactions with both caspofungin and amphotericin B against *C. albicans* but showed a complex interaction with fluconazole. The FICI for compound B and fluconazole is 8.25 due to the 8-fold increase in fluconazole concentration in the combination; however, the concentration of compound B in at FIC is reduced by 4-fold relative to compound A as a single agent. Interestingly, Edlind et al. also observed that cetylpyridinium chloride showed antagonism with fluconazole and induced expression of the efflux pumps *CDR1* and *CDR2* (13).

In contrast to the antagonistic interaction of compound B with fluconazole in *C. albicans*, voriconazole, the gold standard therapy for aspergillosis, showed an additive interaction. Indeed, the activity of compound B was improved 8-fold in combination with voriconazole at ½ MIC. These data indicate that the induction of relative azole resistance in *C. albicans* by B does not occur with *A. fumigatus*. Efflux pump mediated resistance to azoles is not as prevalent in *A. fumigatus* compared to *C. albicans* and, thus, the mechanisms of efflux pump induction and function are not as well defined.

### KAPs are active against *Candida* biofilms and *A. fumigatus* hyphae

In general, pyridinium cations are too toxic for systemic administration because many examples of this class directly lyse red blood cells (15). Consistent with those expectations, compound A and B cause significant red blood cell lysis (LD_50_ A: 26.9 µg/mL; B: 78.6 µg/mL; and C > 1000 µg/mL, Fig. S1) while compound C, which has no antifungal activity also had no activity against red blood cells. This observation provides implicit support for the membrane as an important target of these molecules. Both A and B also showed toxicity against the human cell line HegG2 with LD_50_ of 9 µg/mL and 20 µg/mL using cell lysis and metabolic activity assays, respectively (Fig. S1). Consequently, the KAPs, like other quaternary nitrogen anti-infectives, are likely to be suitable for topical or other non-systemic applications and not feasible for systemic therapy.

As discussed above, many anti-infectives have dramatically reduced activity against biofilm phase organisms compared to the planktonic growth phase of standard CLSI activity assays. To test the activity of the compounds against fungal biofilms, we generated a 24 hr biofilm of *Candida albicans* and then treated with a dilution series of the compounds for an additional 24 hr. The metabolic activity of the biofilms was then assayed using the standard XXT reduction assay (19). Consistent with planktonic results, compound A was active (Fig. 2A, IC_50_ = 18.4 µg/mL) while C was not. The biofilm activity of compound A was reduced by 4-fold relative to planktonic growth (MIC 4 µg/mL, Table 1). Interestingly, the activity of compound A was slightly higher against *C. auris* biofilms (Fig. 2B IC_50_ = 5 µg/mL) relative to planktonic growth (MIC 8 µg/mL, Table 1).

**Figure 2.**
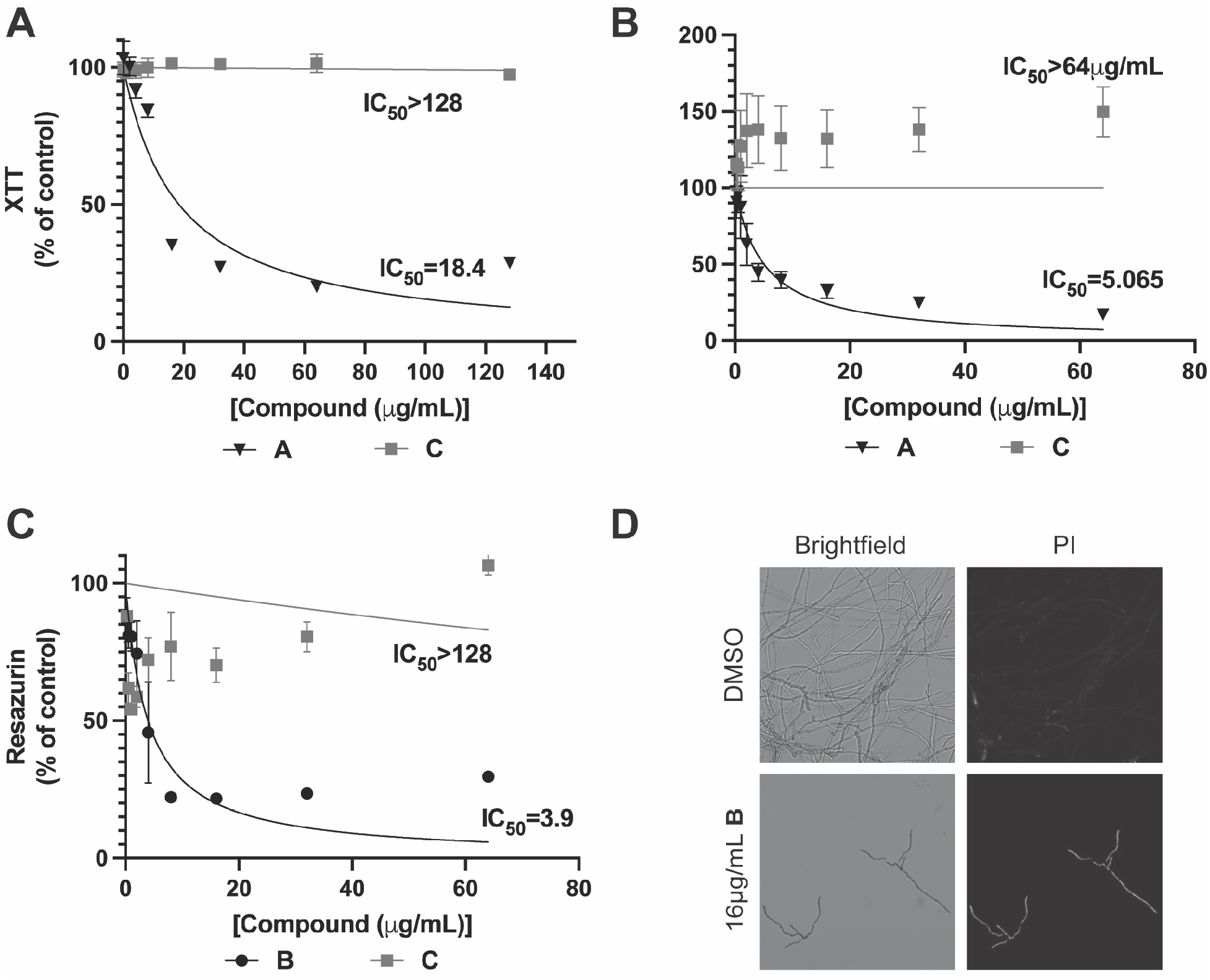
KAPs are active against fungal biofilms. A. Metabolic activity of *A. fumigatus* CEA10 biofilms (measured using resazurin) treated with increasing concentrations B or C for 24 hours. B. Propidium Iodide (PI) staining of *A. fumigatus* CEA10 hyphal cells treated with 16µg/mL B for 24 hours. Images were acquired using the same exposure and magnification for all samples. Representative images from two independent experiments. Metabolic activity of established C. *albicans* SC5314 (**C**) or *C. auris* 0381 (**D**) biofilms treated with **A** or **C** for 24 hours. For all graphs, data represent mean and SEM of three biological replicates and are normalized to untreated controls. IC_50_ curves and values were calculated using GraphPad Prism 9.

CLSI testing of antifungal activity of moulds is based on inoculation with conidia and, therefore, measures inhibition of germination (20). For example, voriconazole is the gold standard for treatment of pulmonary infection and inhibits germination. The most important *A. fumigatus* infection that requires non-systemic therapy is fungal keratitis (21). At the time of clinical presentation, *A. fumigatus* is exclusively in the hyphal form. We, therefore, were interested to determine if compound A or B was active against *A. fumigatus* hyphae. *A. fumigatus* CEA10 was incubated for 24 hr prior to exposure to a dilution series of compound B or C for an additional 24 hr. The metabolic activity of the cultures was determined using resazurin as previously described (17). The IC_50_ (3.9 µg/mL, Fig. 2C) of compound B against this strain was slightly lower than its MIC (8 µg/mL, Table 1) determined by CLSI methods. To further characterize the mode of action of the KAPs against *A. fumigatus* hyphae, we treated CEA10 with compound B for 24 hr and then stained the cells with propidium iodide, a dye that is excluded from cells with intact membranes (Fig. 2D). Whereas cells treated with DMSO control showed essentially no staining, B-treated cells showed uniform uptake of dye, indicating loss of membrane integrity. Taken together, these data indicate that KAPs directly kill hyphal stage *A. fumigatus*.

### KAP A is efficacious in an in vivo model of *C. albicans* venous catheter infection

One potential approach to managing central venous catheter infections is to treat the lumen of the catheter with an anti-infective solution that is locked within the catheter and not introduced into the patient (22). This strategy is most often applied to catheters infected with bacteria but has been proposed for fungal infections as well (5, 6). To test the efficacy of KAPs in this setting, a rat model of *C. albicans* venous catheter infection was employed (23). The catheter was infected with *C. albicans* SC5314 and, on post-infection day 1, treated with A (8 µg/mL solution within catheter) or vehicle. After 24 hr, the catheter was removed and processed for fungal burden and scanning electron microscopy. Compound A-treated catheters showed > 1 log_10_ reduction in fungal burden relative to vehicle-treated catheters (Fig. 3A). Consistent with these results, the treated catheters showed dramatic in the extent of remaining biofilm. Thus, compound A is able to disrupt a pre-formed *C. albicans* biofilm in vitro and in vivo.

**Figure 3.**
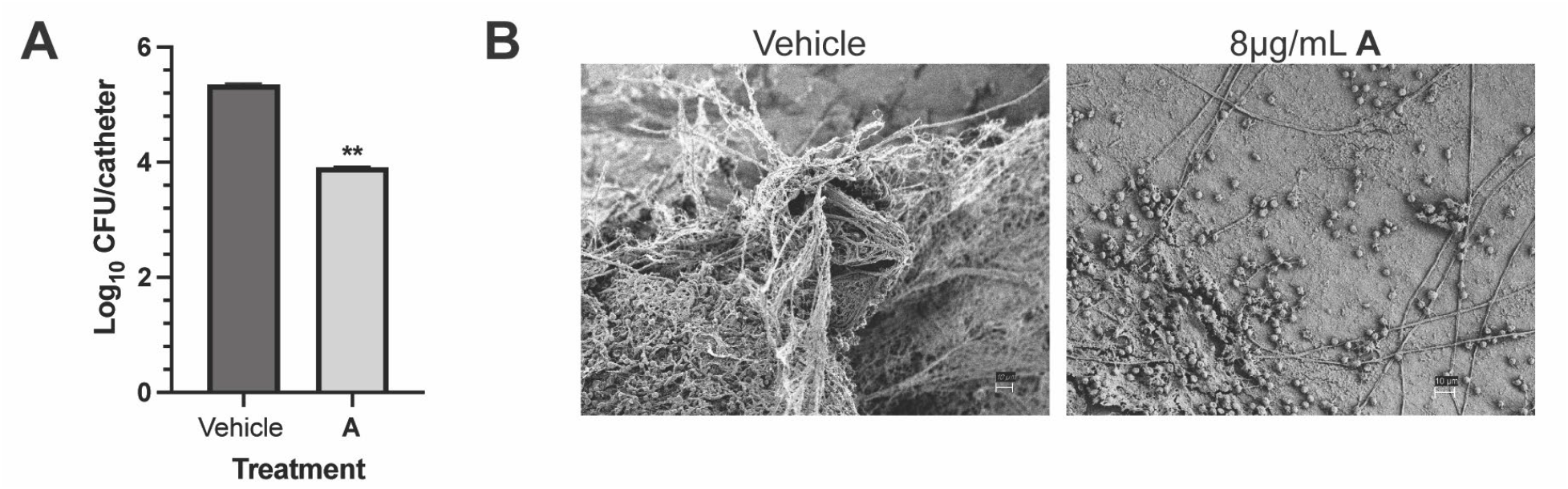
KAPs are efficacious in a rat model of *C. albicans* infection of a vascular catheter. **A**. Colony forming units (CFU) of *C. albicans* from central venous catheters places in rats and treated with vehicle or **A** (8 µg/mL). Fungal burden data are shown as the mean and SEM of log-transformed CFU values; n=3 rats per group. **p<0.0001 by unpaired t-test of log-transformed values. **B**. Scanning electron micrographs of catheters treated with vehicle or 8 µg/mL **A**. Scale bar = 10 µm.

### KAP A reduces *C. auris* colonization of skin in an ex vivo porcine model

The ability of *C. auris* to colonize human skin is likely associated with its persistence in infected patients and its ability to transmit from person-to-person (7). Because compound A showed comparable activity against *C. auris* during planktonic and inanimate-substrate biofilm growth, we tested its activity in an ex vivo porcine model of *C. auris* skin colonization (24). Consistent with these in vitro data, compound A reduced the fungal burden of the colonized porcine skin by ∼1 log_10_ at a concentration of 8 µg/mL (Fig. 4). The fungal burden was not reduced further at 16 µg/mL of compound A, suggesting that absorption/adsorption or physicochemical factors may limit efficacy. The activity of compound A was greater than that of 2% chlorhexidine (∼0.5 log_10_ reduction, ref. 8) and comparable to the synergistic activity of 2% chlorhexidine/70% isopropanol (1.0 log_10_ reduction, ref. 8). We note that, due to limitations in compound A availability, that the data for skin treated with compound A was collected 24 hr after a single treatment while the studies with chlorhexidine and 70% isopropanol were based on a 72 hr experiment with three daily applications of study compounds. These experimental distinctions notwithstanding, compound A shows promising activity against *C. auris* and is likely at least as effective, and possibly more effective, than 2% chlorhexidine.

**Figure 4.**
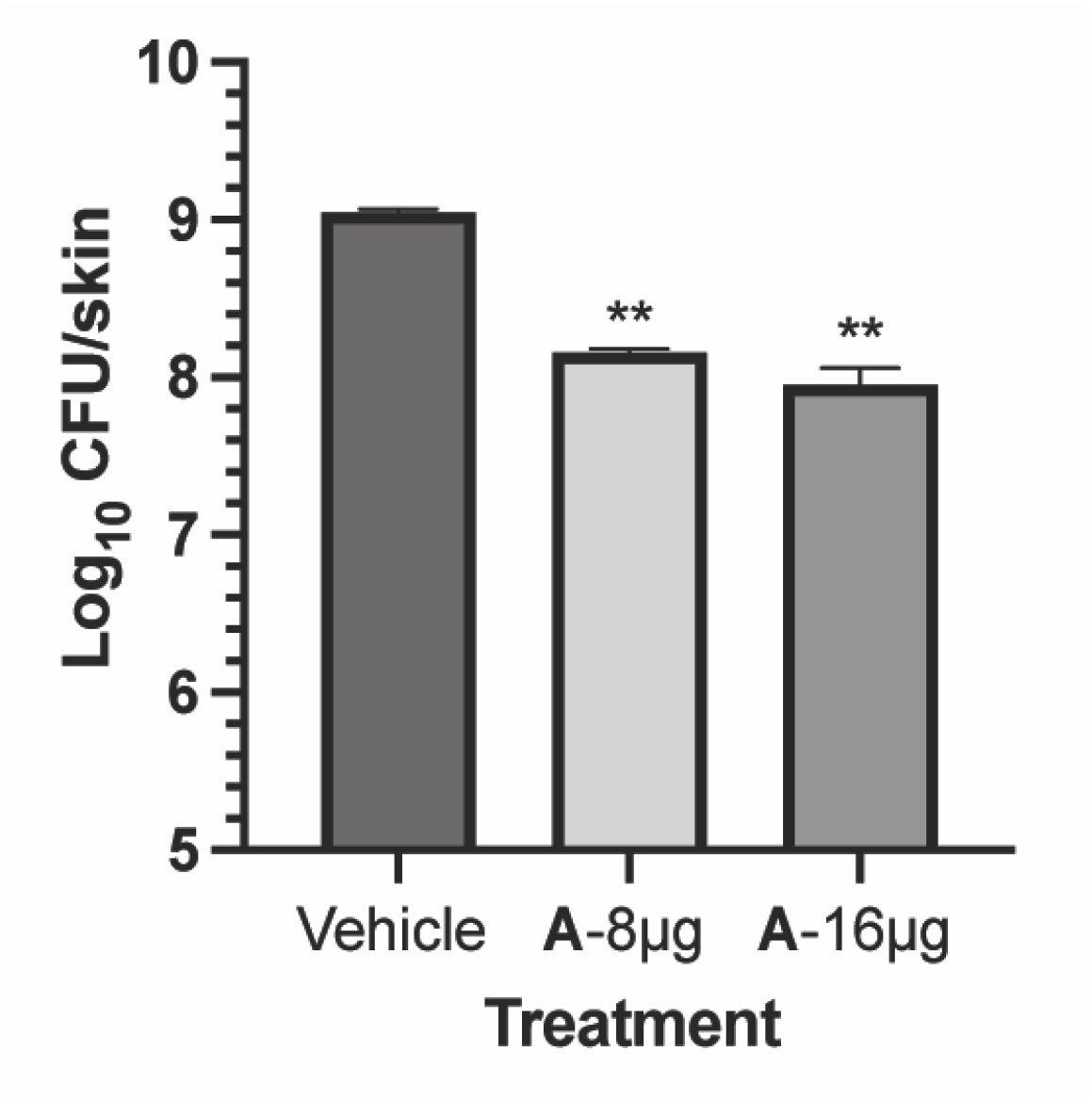
KAP A reduces *C. auris* colonization of skin in an ex vivo porcine model. CFUs of *C. auris* on porcine skin treated with vehicle or **A** (8 µg/mL or 16 µg/mL) for 24 h. Data represent mean and SEM of log-transformed values; n=12 per group. **p<0.0001 by unpaired t-test of log-transformed values.

## Discussion

Although new therapeutic options for systemic therapy of fungal infections are sorely needed (2), agents with promise in the treatment of other types of fungal infections, particularly those involving biofilm and biofilm-like growth phases, would also be quite valuable (3,4). Here, we have characterized the activity of a structurally novel series of pyridinium cation-based antifungal molecules and demonstrate their potential for the treatment of non-systemic fungal infections. Compounds A and B show comparable or improved activity relative to other cationic nitrogen-based anti-infectives/anti-septics such as chlorhexidine and cetylpyridinium chloride against fungal pathogens (8, 13, 14). In addition, A and B have similar activity to the pyridinium-based drug pyrvinium pamoate against *C. auris* against in vivo biofilms (14), further supporting the utility of this class of molecule against this important emerging and drug-resistant pathogen.

Our in vivo and ex vivo experiments provide compelling proof-of-principle data for the potential use of these scaffolds in the setting of antifungal lock therapy for intravascular catheters and as possible disinfectants for *C. auris* skin colonization. With respect to the latter, compound A is more active than the widely studied chlorhexidine (8) which was shown to have activity in a mouse model of colonization (25). Since these compounds are variations on previously validated chemical structures with application to topical and other non-systemic uses, a strong premise exists for their potential development. Additional potential applications for which our data provide support are oropharyngeal candidiasis and fungal keratitis. The latter infection is a globally important cause of blindness for which there is no generally effective medical therapy (21). Compounds A and B have reasonable activity against the two most common etiologic agents of fungal keratitis: *A fumigatus* and *Fusarium* spp (Table 1). The direct activity against hypha is an important feature of this series but additional optimization will be needed to improve the consistency of activity and therapeutic index to become useful for fungal keratitis.

The mechanism of action for quaternary nitrogen-based antifungals such as the KAPs is likely to involve, at least in part, the direct disruption of membrane structures (17). Recent work by Sim et al. suggested that pyrvinium pamoate interferes with mitochondrial function (14) and mode of action studies of the cationic amidine antifungal candidate T-2307 as well as other cationic ammonium antifungals also showed that mitochondrial disruption contributes to their activity (12, 26). A hallmark of mitochondrial disruption in *C. albicans* is an inability to grow on non-fermentable carbon sources such as glycerol. Indeed, the MIC of T-2307 decreases dramatically when the cells are grown on 2% glycerol compared to 2% dextrose (26). We, however, did not detect a change in MIC for compounds A or B when tested in the same medium with glycerol instead of glucose as carbon source, indicating that mitochondrial disruption is unlikely to be a major mode of action (data not shown). Attempts to generate resistant mutants by serial passaging were also unsuccessful which further suggests either multiple targets or non-protein targets such as the membrane are likely to contribute to the antifungal activity of the KAPs.

It is interesting that compound A that, like cetylpyridinium chloride, is antagonistic with fluconazole in *C. albicans* in checkerboard assays. Edlind et al. showed that cetylpyrdinium chloride induces the expression of the efflux pumps *CDR1* and *CDR2* but that deletion of these genes has no effect on its activity (13). Compounds A and B are active as reference strains against fluconazole-resistant clinical strains shown to have increased expression of efflux pumps (Table 1, ref. 18). Thus, it seems that both types of pyridinium based antifungal molecules alter expression of fluconazole pumps without being susceptible to their effects. As such, these observations imply that they have similar mechanisms of action that are best attributed to membrane disruption.

In summary, we provide in vitro and in vivo data suggesting that the KAP-type pyridinium cation molecules may be promising new agents for the non-systemic treatment of fungal infections.

## Materials and methods

### Strains, media, reagents and instrumentation

All yeast strains were maintained on YPD from 25% glycerol stocks stored at −80°C. All *A. fumigatus* strains were maintained on glucose minimal media (GMM; (33) from 25% glycerol stocks stored at −80°C. *A. fumigatus* CFF clinical isolates were received from Dr. Robert Cramer (Dartmouth College), SPF98 was received from Dr. W. Scott Moye-Rowley (University of Iowa), *C. neoformans* DUMC clinical isolate series were received from Dr. John Perfect (Duke University). Reagents were purchased as at least reagent grade from Aldrich, Acros or Alfa Aesar unless otherwise specified and used without further purification. Solvents were purchased from Fischer Scientific (Pittsburgh, PA) and were either used as purchased or redistilled with an appropriate drying agent. Compounds used for structure−activity studies were synthesized according to methods described below, and all the compounds were identified to be least 95% pure using HPLC. Analytical TLC was performed using precoated Silica G TLC Plates, w/UV254, aluminum backed, purchased from Sorbtech (Norcross, GA) and visualized using UV light. Flash chromatography was carried out using with a Biotage Isolera One (Charlotte, NC) system using the specified solvent. Microwave reactions were performed using Biotage Initiator+ (Charlotte, NC). Purity analysis were performed on an Agilent 1100 HPLC utilizing a C-18 column (Waters Nova-Pak; 3.9 × 100 mm) with the following method: Solvent A = H_2_O (0.1% TFA), Solvent B = Acetonitrile; 0 to 20 min, (10 to 90% B), 20 to 25 min (90 to 10% B); detection was set at two wavelengths (245 and 280 nm). Purity of all final compounds was above 95%. All final compounds were analyzed by high resolution MS (HRMS) using a Bruker Maxis Plus Quadrupole Time-of-Flight (QTOF or QqTOF). ^1^H and ^13^C NMR were recorded on either a BrukerAvance III 500 outfitted with a 5mm BBFO Z-gradient probe or a Avance III 300 instrument is equipped with a 5 mm BBFO probe. The chemical shifts are expressed in parts per million (ppm) using suitable deuterated NMR solvents.

### Dichlorodiphenylmethane (1)

To a round flask was weighed benzophenone (1eq) and PCl_5_ (1.5eq). The reaction was refluxed for 2 h at 150°C. After 2 hours, the reaction was cooled, 40 mL of DCM was added to the resultant solution followed by transferring to 250 mL Erlenmeyer flask. The resultant solution was cooled to 0°C and 40 mL of distilled water was added. The resultant solution was transferred to a separatory funnel, the DCM layer was separated, washed twice with 40 mL distilled water, and dried with anhydrous sodium sulfate. The DCM solvent was evaporated in rotary evaporator to give corresponding dichloride. The crude product was immediately used for the next step without purification.

### 2-chloro-1-(2,2-diphenylbenzo[d][1,3]dioxol-5-yl)ethenone (2)

To a round flask was weighed dichlorodiphenylmethane (1 g, 4.2 mmol) and 2-chloro-1-(3, 4-dihydroxyphenyl) ethanone (0.79 g, 4.2 mmol) under nitrogen. The reaction was refluxed at 180°C for 30mins under nitrogen. After 30 mins. The crude product was transferred to silica gel chromatography with linear gradient of 2 - 10% (EtOAc/Hexane). The fractions corresponding to the desired product were collected and solvent evaporated using rotatory evaporator to give yellow viscous liquid as the desired product which turned to a solid upon dryness. Yield; 0.7 g, 47%, ^1^H NMR (500 MHz, CDCl_3_) δ 7.69 (m, 4H), 7.61-7.57 (m, 2H), 7.44 (m, 6H), 6.98 (d, *J* = 7.9 Hz, 1H), 4.63 (s, 2H).^13^C NMR (75 MHz, CDCl_3_), 189.26, 152.05, 148.05, 139.46, 129.56, 129.01, 128.41, 125.14, 118.58, 108.52, 108.38, 45.84

### 4-acetamido-1-(2-(2,2-diphenylbenzo[d][1,3]dioxol-5-yl)-2-oxoethyl)pyridin-1-ium chloride (3)

To a round flask containing 2-chloro-1-(2,2-diphenylbenzo[d][1,3]dioxol-5-yl)ethenone (0.34 0.97 mmol) dissolved in acetonitrile (10 mL) was weighed the 4-aminopyridine (0.09 g, 0.97 mmol). The reaction was refluxed for 2hrs at 100°C in which precipitate was formed. After 2 hrs, the solvent was evaporated using rotatory evaporator and the gummy solid precipitated with 10% Hexane in DCM, filtered and dried to give a powdered solid product in form of the chloride salt. HPLC purity; 98%. Yield;0.38 g, 88%, ^1^H NMR (300 MHz, DMSO) δ 12.12 (s, 1H), 8.65 (d, *J* = 6.9 Hz, 2H), 8.22 (d, *J* = 6.9 Hz, 2H), 7.74 (d, *J* = 8.3 Hz, 2H), 7.68 (s, 1H), 7.57-7.55 (m, 4H), 7.48-7.32 (m 6H), 7.32 (d, *J* = 8.2 Hz, 1H), 6.21 (s, 2H), 2.27 (s, 3H).^13^C NMR (75 MHz, DMSO), 189.88, 171.71, 152.87, 151.87, 147.59, 146.99, 139.30, 130.21, 129.18, 129.00, 126.25, 125.70, 118.36, 114.77, 109.56, 108.65, 64.75, 25.06, HRMS for [M+H]^+^ calculated; 451.1652, found; 451.1658

### Planktonic growth phase antifungal activity assays

Minimum inhibitory concentrations were determined using CLSI guidelines (16, 20). All yeasts were cultured overnight in 3 mL YPD at 30°C, then washed twice in sterile PBS. Two-fold serial dilutions of each compound were prepared in RMPI+MOPS pH 7 (Gibco RPMI 1640 with L-glutamine [11875-093] and 0.165M MOPS), then 1 × 10^3^ cells were added per well. Plates were incubated at 37°C for 24 h (*C. albicans* and *S. cerevisiae*) or 72 h (*C. neoformans*). For *A. fumigatus* and *F. oxysporum* wells were inoculated with 1.25 × 10^4^ conidia. Plates were incubated at 37°C for 48 h for *A. fumigatus* and 72 h for *F. oxysporum*.

### Biofilm phase antifungal activity assays

These assays were performed as previously described (17, 19). *C. albicans* and *C. auris*, overnight cultures were washed in sterile PBS and adjusted to 1×10^6^ CFU/mL in RPMI+MOPS pH 7. 100µL cells were added to each well and incubated at 37°C for 24 h. A two-fold dilution series of the compounds was prepared in RPMI + MOPS with equal vehicle concentrations. Media was removed and biofilms were gently washed with PBS, then 200µL of each drug dilution was added to wells and plates were incubated at 37°C for an additional 24 h. Media was removed and biofilms were gently washed with PBS, then 100 µL of XTT solution (0.83 mg/mL XTT sodium salt [Sigma, Cat# X4626] + 32 µg/mL PMS [Sigma, Cat# P9625] in PBS) was added to each well. Plates were incubated at 37°C for 30 min, then the absorbance was measured at 490 nm.

### Red blood cell lysis assay

Hemolysis assays were performed with defibrinated sheep’s blood (Lampire, Cat# 7239001). Blood was washed three times with PBS then resuspended to ∼50% hematocrit in PBS. A two-fold dilution series of the compounds in 200µL PBS with equal DMSO concentration across the series, then red blood cells were added with a final concentration of 2% hematocrit. Cells were incubated at RT for 2 h in a v-bottom microtiter plate, then plates were spun down and supernatant was transferred to a flat bottom microtiter plate for absorbance measurement at 570 nm. Compounds were tested in technical triplicate in at least two independent assays performed on different days.

### Mammalian cell culture toxicity assay

HepG2 cells (ATCC, #) were maintained and cultured in HepG2 media (DMEM (Gibco, Cat#11965-092) with 5% FBS, 20 mM Glutamine, and Penicillin/Streptomycin) at 37°C with 5% CO_2_. Cells were seeded in 96-well plates at a density of 1.25 × 10^4^ cells/well and incubated overnight at 37°C with 5% CO_2_, then media was removed and replaced with media containing a two-fold dilution series of compounds with equal DMSO concentrations across all wells including no compound controls. Cells were incubated for an additional 24 h. Supernatant was removed and used to quantify lactate dehydrogenase release using the CyQuant LDH assay kit (Invitrogen, Cat#C20300) following the manufacturer’s directions. LDH signal was normalized to a max lysis control. The remaining cells were then used to quantify cellular metabolism by XTT. Briefly, cells were washed, then 0.9 mg/mL XTT (Sigma, CAT#X4626) + 320 µg/mL PMS (Phenothiazine methosulfate; Sigma, Cat# P9625) in HepG2 media was added to each well. Plates were incubated at 37°C, with 5% CO_2_ for 2 h, then absorbance was measured at 490 nm.

### Rat model of *Candida albicans* vascular catheter infection

*C. albicans* biofilm growth during infection of implanted medical devices was measured using an external jugular-vein, rat-catheter infection model (23). Briefly, a 1 × 10^6^ cells/ml inoculum for each strain or strain combination was allowed to grow on an internal jugular catheter placed in a pathogen-free female rat (16-week old, 400 g) for 24 h. After this period, the catheter volumes were removed and the catheters were flushed with 0.9% NaCl. The biofilms were dislodged by sonication and vortexing. Viable cell counts were determined by dilution plating. Three replicates were performed for each strain.

Scanning electron microscopy of catheter biofilms. After a 24 h biofilm formation phase, the devices were removed, sectioned to expose the intraluminal surface, and processed for SEM imaging. Briefly, one milliliter fixative (4% formaldehyde and 1% glutaraldehyde in PBS) was added to each catheter tube and tubes were fixed at 4 °C overnight. Catheters were then washed with PBS prior to incubation in 1% OsO4 for 30 min. Samples were then serially dehydrated in ethanol (30–100%). Critical point drying was used to completely dehydrate the samples prior to palladium–gold coating. Samples were imaged on a SEM LEO 1530, with Adobe Photoshop 2022 (v. 23.2.2) used for image compilation.

### Porcine skin model of *Candida auris* colonization

The collection of porcine skin samples was conducted under protocols approved by the University of Wisconsin–Madison Institutional Animal Care and Use Committee in accordance with published National Institutes of Health (NIH) and United States Department of Agriculture (USDA) guidelines. Excised skin was cleaned and shaved as described previously (24). Full-thickness samples were obtained by 12 mm punch and placed in 12-well plates containing 3 mL Dulbecco’s Modified Eagle Medium (DMEM) (Lonza, Walkersville, MD, USA) supplemented with 10% fetal bovine serum (FBS) (Atlanta biologicals, Lawrenceville, GA, USA), penicillin (1,000 U/mL), and streptomycin (1 mg/mL) (Corning, Manassas, VA, USA) for 6 hours. Tissues were rinsed in DBPS and moved to semi-solid media (6:4 ratio of 1% agarose (BIO-RAD, Hercules, CA, USA) in DPBS:DMEM with 10% FBS). Paraffin wax was applied around the edge of the skin and 10 μL *C. auris* suspended in synthetic sweat medium at 10^7^ cells/mL was applied. Samples were incubated at 24 h and compound A (8 µg/mL or 16 µg/mL) in synthetic sweat medium was applied. After 24 h a sterile swab was used to remove compound from skin surface. Samples were vortexed in DPBS and plated on YPD + chloramphenicol to assess viable burden.

## Acknowledgements

This work was funded in part by NIH grants: R21AI164578 and R01AI161973 (DJK); F32AI145160 (SRB), and R01AI073289 (DRA). Purchase of the NMR spectrometer used to obtain results included in this publication was supported by the National Science Foundation under the MRI award CHE-2117776. The authors thank Steven Riesinger and Tsvetelina Lazarova (Med Chem Partners) for additional medicinal chemistry support of this project.

## Figure Legends

**Figure S1.**
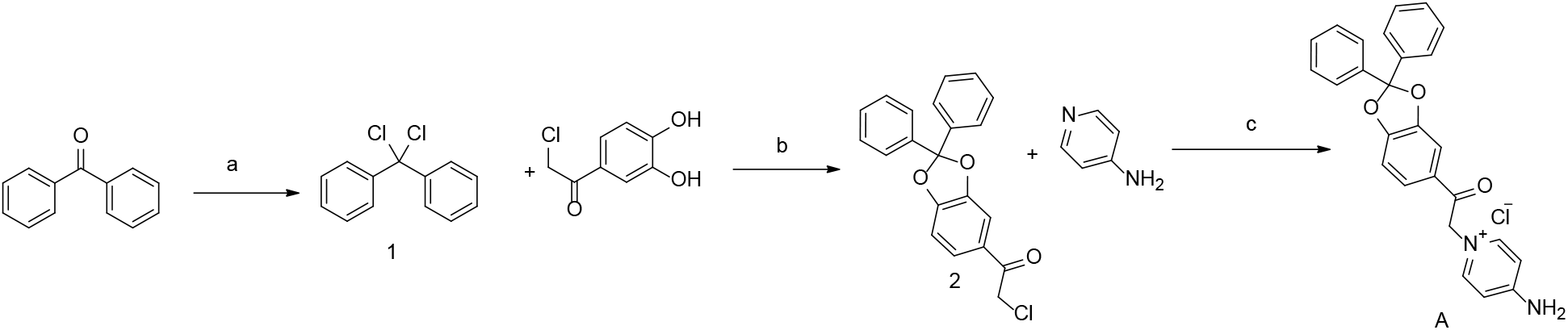
Chemical synthesis of compound. **A**. Reaction conditions: (a) PCl_5_ (1.5 eq), 2 h, reflux 150°(b) 30 min, 180°C, reflux (c) 2 h, 100°C acetonitrile reflux.

**Figure S2.**
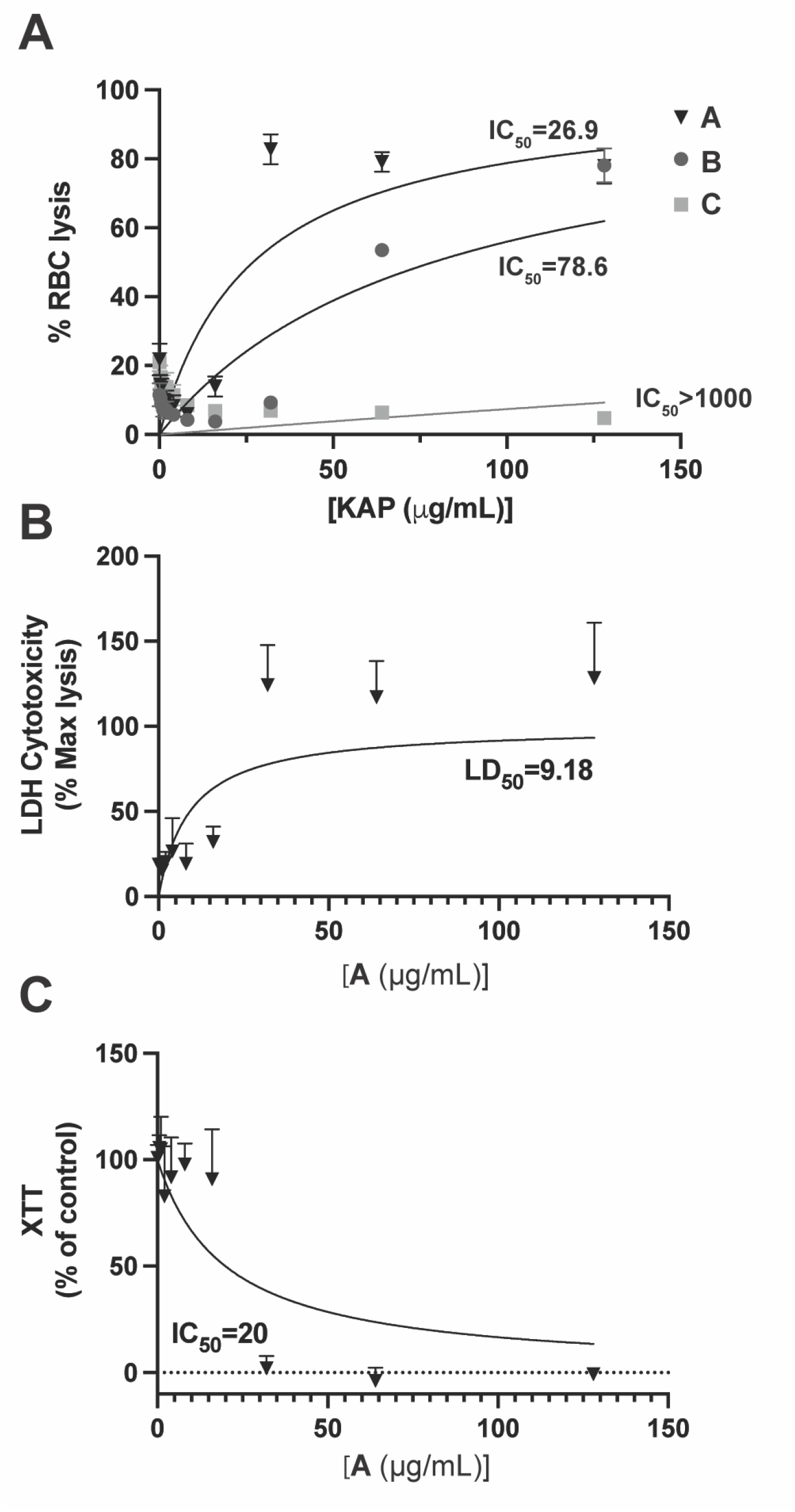
Red blood cell and mammalian cell culture toxicity of KAP compounds. **A**. Hemolysis of commercial red blood cells (RBC) treated for 2 h with increasing concentrations of KAPs normalized to max lysis by triton-x. Mean and SD of technical triplicates. Representative data from three independent experiments performed on different days are shown. Toxicity against HepG2 cells using LDH release (**B**) and XTT (**C**) assays. Cells were treated with the indicated concentration series of **A** for 24 h. Mean and SD of technical triplicates. Data are representative of two independent experiments performed on different days.

